# Non-random polymorphisms near transposable elements facilitate evolution across the genome of *Arabidopsis thaliana*

**DOI:** 10.1101/2023.10.24.563748

**Authors:** H. De Kort, Y. Ganseman, D. Deforce, F. Van Nieuwerburgh

## Abstract

Evolution is a key characteristic of all life on earth, and is achieved mainly through mutation and recombination. To facilitate evolutionary rates in the absence of recombination, increased mutagenesis could provide essential opportunities for evolution. Here, we harness the genomes of 973 wild accessions of *Arabidopsis thaliana*, a self-fertilizing species characterized by reduced recombination rates, to study the role of transposable elements (TEs) as mutagenic drivers of non-random genetic variation. We found that multiple TE superfamilies accumulate large numbers of genetic variants in gene-rich regions of the genome. Moreover, TEs were enriched in genes underpinning key molecular processes, including fertilization and mRNA cis-splicing, suggesting that TEs generate genetic variation that is fundamental to organismal functioning and reproduction. An excess of common genetic variants (maf>0.4) flanking TEs, despite the extreme degree of self-fertilization, further corroborates the notion that TEs can generate substantial evolutionary potential in the absence of outcrossing. Nevertheless, while other studies point to TE mobilization as a strategy to facilitate adaptive evolution, we find that only a fraction (4.8%) of TE families was linked to genetic variants involved in climate adaptation to a significantly larger extent than background genomic variation, possibly because the high mutation rates near TEs dilute adaptive signals. Overall, all TE superfamilies but one (Gypsy) were significantly associated with evolutionary processes through their association with genes, functional molecular processes, flanking common variants and/or climate adaptation. We conclude that non-random mutations flanking TEs provide substantial evolutionary potential in a self-fertilizing species.

## Background

Mutation and recombination provide the molecular variation upon which evolution is based, and thus ensure long-term organismal functioning and population persistence. Understanding variation in mutation and recombination rates is thus fundamental to evolutionary theory. Within individuals, mutation rates have been demonstrated to vary across the genome, with transposable elements (TEs) being tightly linked to mutation peaks (Schrader et al. 2014; Lu et al. 2017; Schrader and Schmitz 2019; Habig et al. 2021a). TEs are mutagenic, self-replicating sequences that are common across most plant and animal genomes, and have been shown to preferentially integrate in gene-rich regions where they can be co-opted to fine-tune gene expression and organismal functioning (McClintock 1984; Stapley et al. 2015; Lu et al. 2017; Sultana et al. 2017a; Baduel et al. 2021). As a consequence, evolution is not merely the result of natural selection acting on random mutations, but can, in theory, be directed by molecular mechanisms (Monroe et al. 2022). However, to what extent TEs guide evolutionary trajectories of flanking genome regions remains poorly explored.

Targeted mutation, i.e. neutral or beneficial mutations occurring preferably in functional genomic regions, may be particularly important to self-fertilizing species with reduced capability of exchanging and reshuffling genetic novelty between genetically variable individuals. Repeated inbreeding results in homozygous organisms with reduced evolutionary potential and intense purifying selection (Noël et al. 2017), unless mutation bias occurs sufficiently to avoid maladaptation and the deleterious impacts of genome-wide homozygosity. An underexplored mechanism potentially driving non-random mutation across the genome involves transposable elements, which can facilitate host evolution (Schrader and Schmitz 2019; Catlin and Josephs 2022; De Kort et al. 2022). Transposable elements (TEs) tend to mobilize following environmental stress (Cappucci et al. 2019), and to subsequently insert into functional genomic regions (Baduel et al. 2021), where they increase flanking methylation and mutation rates (Ossowski et al. 2010; Wicker et al. 2016; Habig et al. 2021b). Therefore, exploring the genomic distribution of TEs and flanking genetic variation could give unique insights into the evolutionary abilities and success of self-fertilizing species challenged by low recombination rates.

The classification, distribution, and abundance of TEs in the self-fertilizing *Arabidopsis thaliana* are well characterized and provide important insights into the functional and evolutionary implications of TE activity and mobilization (Quesneville 2020). Notably, TEs tend to be concentrated in genomic regions implicated in adaptive divergence (Schrader and Schmitz 2019; Baduel et al. 2021), but the possibility that they may be responsible for the mutations underpinning the evolutionary trajectories of self-fertilizing species has not yet been explored. A recent study points to epigenetically-driven mutation bias near gene bodies in *A. thaliana* (Monroe et al. 2022), indirectly suggesting that transposable elements flanking genes cause targeted mutation. Thus, while TEs are typically epigenetically silenced to protect genome integrity and functioning (Kelleher et al. 2020), they may facilitate evolution through their intense association with methylation; a mechanism also referred to as mutation-by-methylation (Habig et al. 2021; De Kort et al. 2022). Particularly in *A. thaliana*, a homozygous species characterized by extreme levels of self-fertilization, such non-random genomic sensitivity to mutation could allow long-term evolution under prolonged self-fertilization and inbreeding.

The ultimate distribution of TEs and flanking genomic variation is driven by host-TE coevolution, where a balance between TE activity and host fitness determines the fate of TEs and their host through purifying and adaptive selection (Kidwell and Lisch 1997; Sultana et al. 2017b). A long history of host-TE coevolution characterized by an accumulation of TEs co-opted in functional genomic regions should thus result in an accumulation of mutations in the same genomic regions. When these mutations are beneficial to the host, their allele frequency should rise unless a lack of outcrossing prevents the spread of new mutations. However, beneficial mutations may become common despite nearly exclusive self-fertilization if they appear repeatedly and non-randomly across the range. While enriched signatures of balancing selection have been observed near TEs in natural inbred populations of *A. lyrata* (De Kort et al. 2022), the genome-wide association of non-random genetic variation with TEs remains obscure. Similarly, signatures of adaptive evolution are expected to accumulate near TEs, corresponding to the high evolutionary rates observed in TE-rich genomic regions (Schrader and Schmitz 2019; Baduel et al. 2021).

Here, we shed light on the genomic and spatial distribution of single nucleotide polymorphisms (SNPs) flanking TEs of all major TE superfamilies and across the Eurasian range of *A. thaliana*. The behavior and functional characteristics of genetic variants near TEs as compared to those arising away from TEs can inform to what extent TEs drive evolution in a homozygous species. Because *A. thaliana* is a self-fertilizing pioneer species able to efficiently colonize ruderal environments across a broad climatic gradient, new mutations are expected to rapidly disappear or shift to fixation due to the combined actions of drift, inbreeding, and selection. Rare alleles thus typically represent relatively recent mutations. We hypothesize that (i) TEs of various TE superfamilies are non-randomly distributed across the genome of *A. thaliana*, (ii) rare genetic variants accumulate rapidly within TEs, corresponding to the high mutation and diversification rates of TEs, and (iii) genetic variants accumulating near TEs provide opportunities for evolution, particularly in functional genomic regions, and are more common across the range than expected for self-fertilizing species.

## Results

### TEs and linked genetic variants are non-randomly distributed across the genome

To address our hypothesis that TEs are non-randomly distributed across the genome, we first explored the tendency of the major *A. thaliana* TE superfamilies to occur near genes (Fig. 1A). We show that with 86% of all TEs organized in blocks consisting of at least two TEs with less than 1 kb in between (Fig. 1A), the large majority of TEs is clustered in the centromeric and pericentromeric regions of the genome (Fig. 1B) and with no marked affinity to genes (Fig. 1C, Table S1). However, the proximity to genes and their tendency to cluster together varies substantially among TE superfamilies, with Gypsy elements typically residing away from genes (Fig. 1C and 1D) and in dense clusters (Fig. 1A), while SINE elements are more evenly distributed (Fig. 1A) and particularly abundant in gene-rich regions (Fig. 1C and 1D, Table 1, Table S1). MuDR elements, which are known for their mutagenic properties (Dupeyron et al. 2019), frequently integrate within or adjacent to genes (Fig. 1C and 1D).

**Fig. 1.**
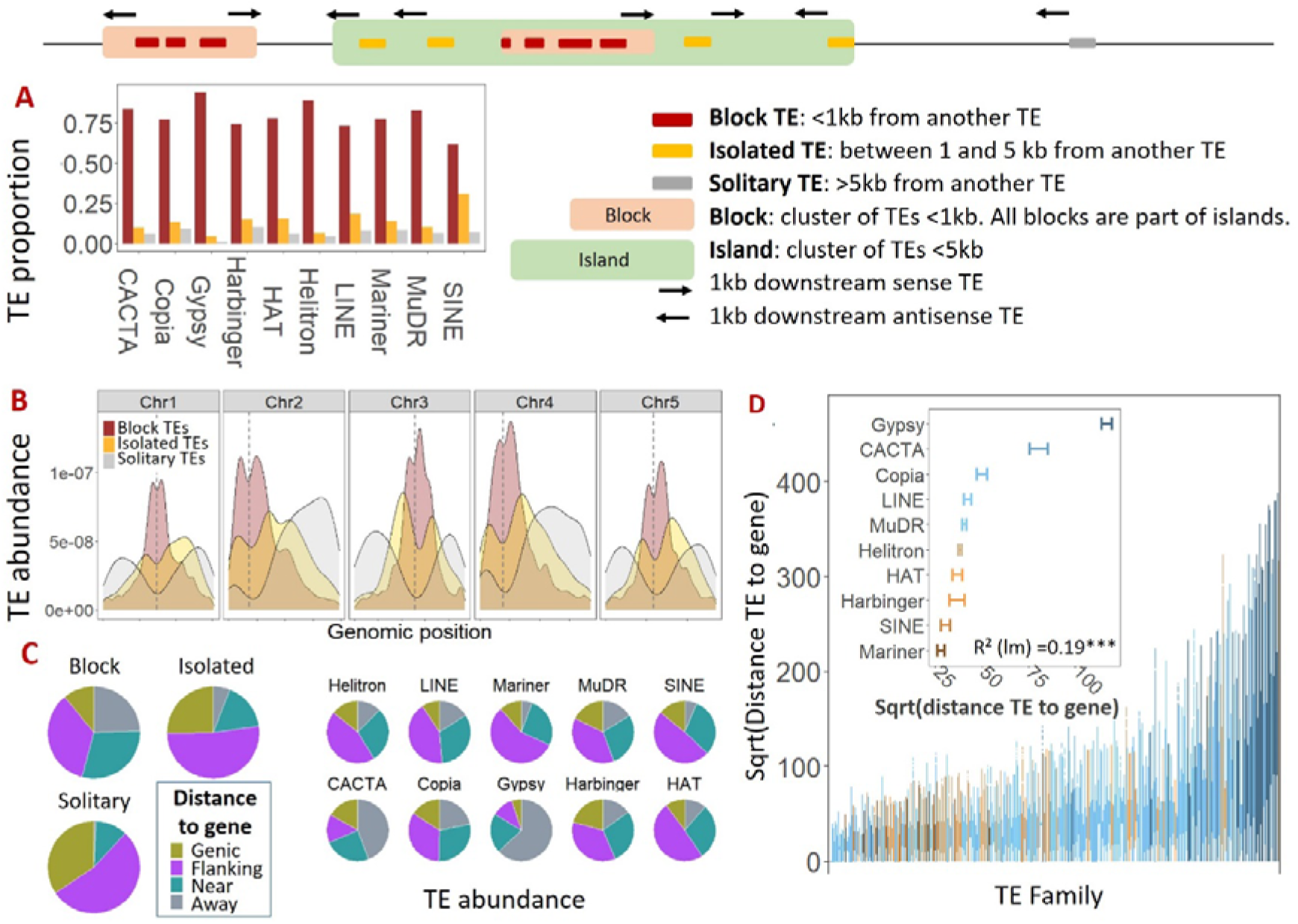
Genomic distribution of transposable elements (TEs). (**A**) Most TEs are within 1 kb of another TE (in TE blocks, see Table S1). All TE blocks are within 5 kb of other TEs (in TE islands). (**B**) The density of TEs in *Arabidopsis thaliana* decreases along the chromosome arms from densely clustered TEs in the centromeres (dashed lines) to solitary TEs near the chromosome ends. (**C**) Highest TE densities can be found adjacent to genes, particularly for solitary TEs. (**C, D**) The proximity to genes varies considerably and significantly among TE superfamilies.

**Table 1.**
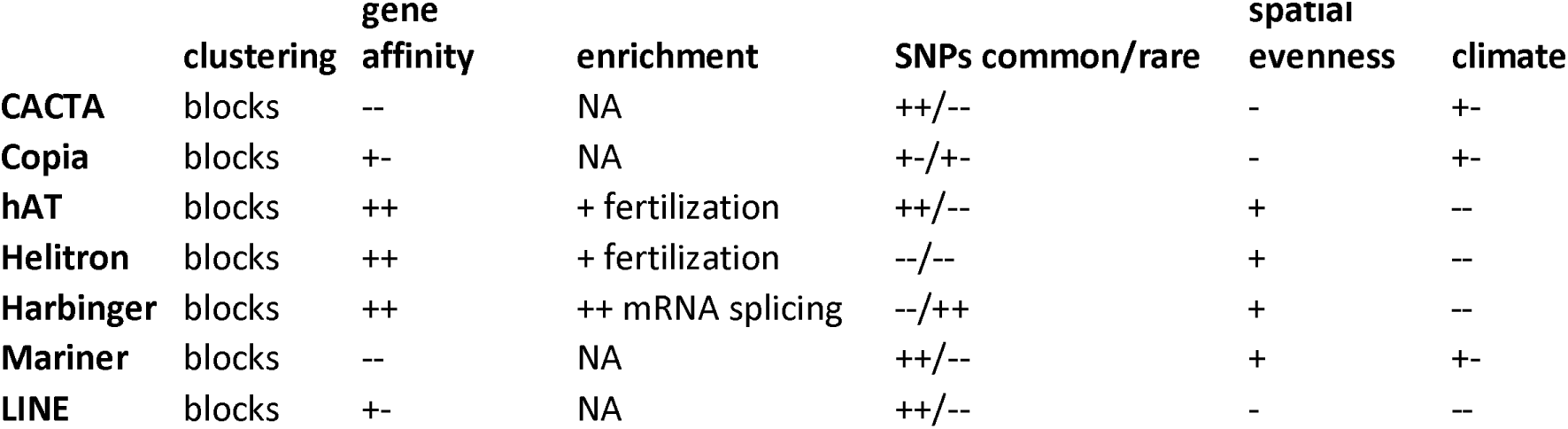
Summary of TE superfamily characteristics. Gene affinity is based on **Fig. 1C**, with less than half, half, or more than half of the TEs within or adjacent to genes (--, +- and ++, resp.). Enrichment is based on **Fig. 2A**, and points to molecular processes significantly enriched for TEs of the respective superfamily (+ or ++). Only the Gypsy family is characterized by a significant underrepresentation of TEs (--). SNPs common/rare is based on Fig. S1 and indicates superfamilies with high (++), moderate (+-) or low (--) proportions of <u>downstream</u> common and rare variants. Spatial evenness and climate are based on **Fig. 6**, indicating the proportion of TE families significantly more (+) or less (-) involved in balancing selection and climate adaptation as compared to background genomic variation.

Gene ontology enrichment analyses revealed that genes free of TEs (TEs at least 5kb from genes) are significantly enriched for chromosome segregation functions, with over double (i.e. 200%) the expected number of genes involved in cell division being located far away from TEs (Fig. 2A, Table S2). Genomic regions away from TEs were significantly enriched for many other processes, including flower development (56% enriched), response to external stimuli (51%), seed development (39%), growth (35%), and root development (21% enriched). For genes overlapping with TEs, Harbinger and MuDR elements manifest the most striking enrichments (Table S2). Specifically, genes containing Harbinger elements are 75-fold enriched for genes performing mRNA cis-splicing (Fig. 2A). Genes containing MuDR elements are 22-fold enriched for genes regulating transcription (Fig. 2A). Genetic variation near Harbinger and MuDR elements are thus likely to associate with nearby gene transcription and expression. Responses to abiotic stimuli are underrepresented by genes adjacent to (<1kb) Gypsy elements (Fig. 2A).

**Fig. 2.**
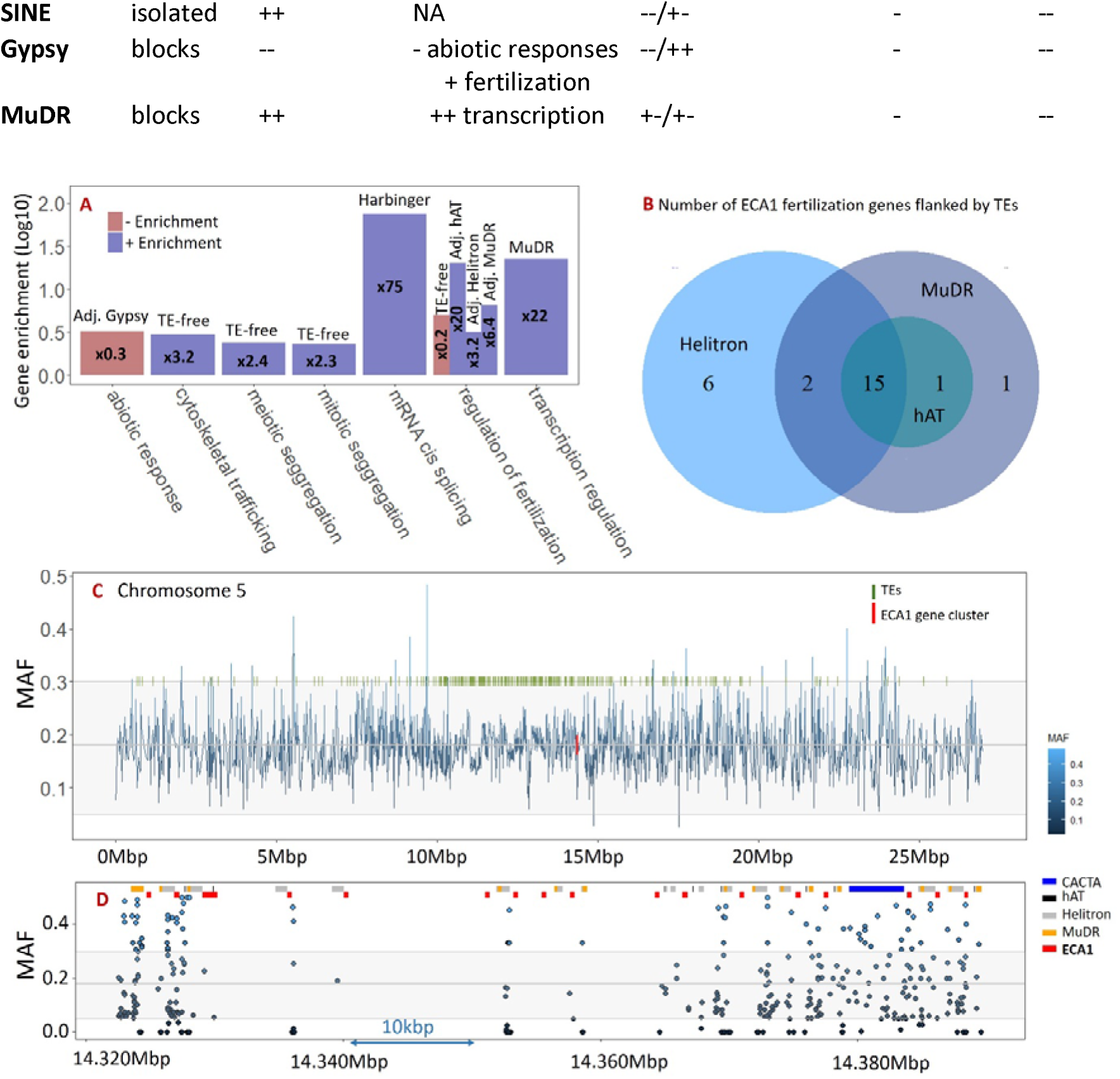
**(A) Enrichment of genes** lacking, containing or adjacent to transposable elements, on a logarithmic scale. Only processes that were at least two-fold enriched (i.e. occurred more than twice than expected) are shown (see Table S2 for a full list of over- and underrepresented processes). Genes free of TEs (> 5kb from TEs) are particularly enriched for cell division processes. **(B)** Fertilization-related genes, ECA1 genes in particular, are notably associated with TEs from three superfamilies (Helitron, MuDR, and hAT). Note that only the three significantly ECA1-enriched TEs are shown; the total number of TEs flanking ECA1 genes adds up to 36 (Table S1). **(C)** A total of 17 ECA1 genes in the ECA1 gene cluster in the pericentromeric region of chromosome 5 was found to be flanked by TEs, and **(D)** the TE-enriched ECA1 gene cluster also contained high SNP densities with varying maf, particularly in the vicinity of MuDR elements (yellow).

Most interestingly, however, is the notable enrichment of fertilization-related genes flanked by elements of the hAT, MuDR, and Helitron superfamilies: At least 36 genes belonging to the unexplored group of ECA1 (Early Culture Abundant 1, of which 124 genes identified in *A. thaliana*) gametogenesis-related cysteine-rich proteins are flanked by transposable elements (Fig. 2B, Table S2). These proteins are involved in double fertilization, a key characteristic of angiosperm reproduction. Half of these genes (N=17) reside within a 100kbp pericentromeric region on chromosome 5 (Fig. 4C and Fig. 4D), where they are associated with high TE densities and highly variable minor allele frequencies (Fig. 4D).

To explore the tendency of TE superfamilies to accumulate genetic variation, we first investigated the distribution of genetic variants (SNPs) extracted from the publicly available 1001 *A. thaliana* genome project (Alonso-Blanco et al. 2016) in relation to nearby TEs. For all 973 Eurasian accessions (Table S3), we extracted all SNPs within and adjacent to TEs (N=500,581 and 328,665 SNPs resp.; maf>0.05). We focused on the 1000bp downstream region to characterize SNPs directly flanking TEs.

We found that solitary TEs that are at least 5 kb from other TEs have similar amounts of nearby genetic variants as TEs clustered in TE blocks (Fig. 3A). However, because solitary TEs reside in gene-rich regions of the genome (Fig. 1C), they predominantly generate genetic variation in functional parts of the genome. Because SNPs downstream of TEs are likely to influence genome functioning and evolutionary trajectories (De Kort et al. 2022), we then studied the tendency of TEs to accumulate downstream SNPs. The SNP density downstream of TEs correlates with the SNP density within TEs, but for every increase in downstream SNP density, the SNP density within TEs increases disproportionally (Fig. 3C, Fig. S2). This bias towards SNP accumulation within TEs is particularly pronounced for TE superfamilies flanking genes, where most TEs are characterized by relatively low downstream SNP accumulation (Fig. 3D). Near genes, purifying selection may thus reduce the number of SNPs downstream of TEs. (Fig. 3D). TEs away from genes, on the other hand, tend to accumulate high numbers of downstream SNPs (Fig. 3D).

**Fig. 3.**
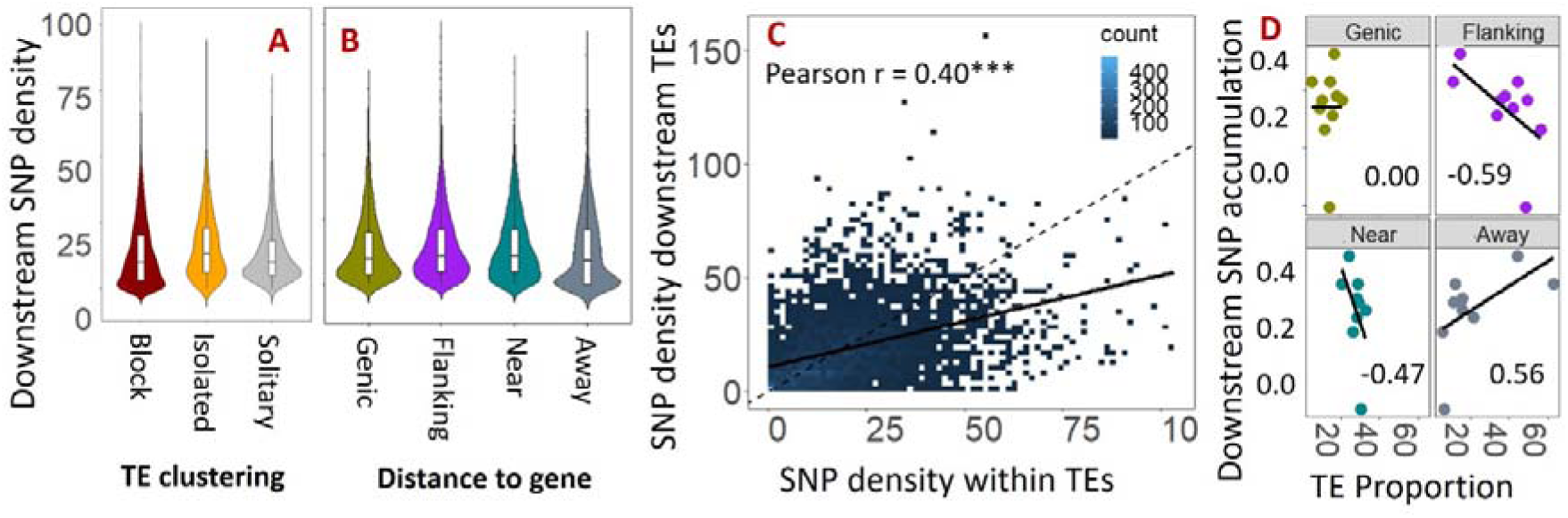
SNP accumulation downstream of TEs. **(A)** The density of SNPs downstream of TEs does not markedly vary between block, isolated and solitary TEs, and (**B**) does not depend on the distance of TEs to the nearest gene. (**C**).The SNP density downstream of TEs correlates with the SNP density within TEs (significant Pearson r), but SNPs accumulate more rapidly within than downstream of TEs. (**D**) Downstream SNP accumulation here refers to the TE superfamily-specific slope of the regression line in panel C, with a steeper slope representing the tendency of variable TEs to also accumulate downstream SNPs. (**D**) TE superfamilies with a high proportion of TEs flanking genes are characterized by low downstream SNP accumulation irrespective of the SNP densities within TEs. On the other hand, TE superfamilies with a high proportion of TEs away from genes are characterized by high downstream SNP accumulation. The outlier point represents the SINE superfamily, which can reach large SNP densities within TEs (Fig. S2), but without a parallel increase in downstream SNP densities, and is also characterized by a relatively high proportion of TEs flanking genes (Fig. 1C and 1D).

### Rare and common variants within and downstream of TEs

To test our second hypothesis that rare genetic variants accumulate rapidly within TEs due to their high mutation and diversification rates, we analyzed the allele frequencies (maf>0.05) of SNPs within and downstream of TEs, and compared them to background SNPs > 5kb away from any TE (N=8654). In congruence with the high mutation and diversification rates of TEs, we found an enrichment of rare genetic variants (maf<0.06) within TEs (Fig. 4A). Thus, while overall SNP densities are higher within TEs than downstream of TEs, these SNPs typically have low frequencies. This is not the case for the genomic regions directly downstream of TEs, where instead, common alleles (maf > 0.45) are enriched relative to background genetic variants (Fig. 4B). Common alleles are unlikely to represent deleterious mutations and instead point to standing genetic variation providing contemporary or cryptic opportunities for natural selection.

**Fig. 4.**
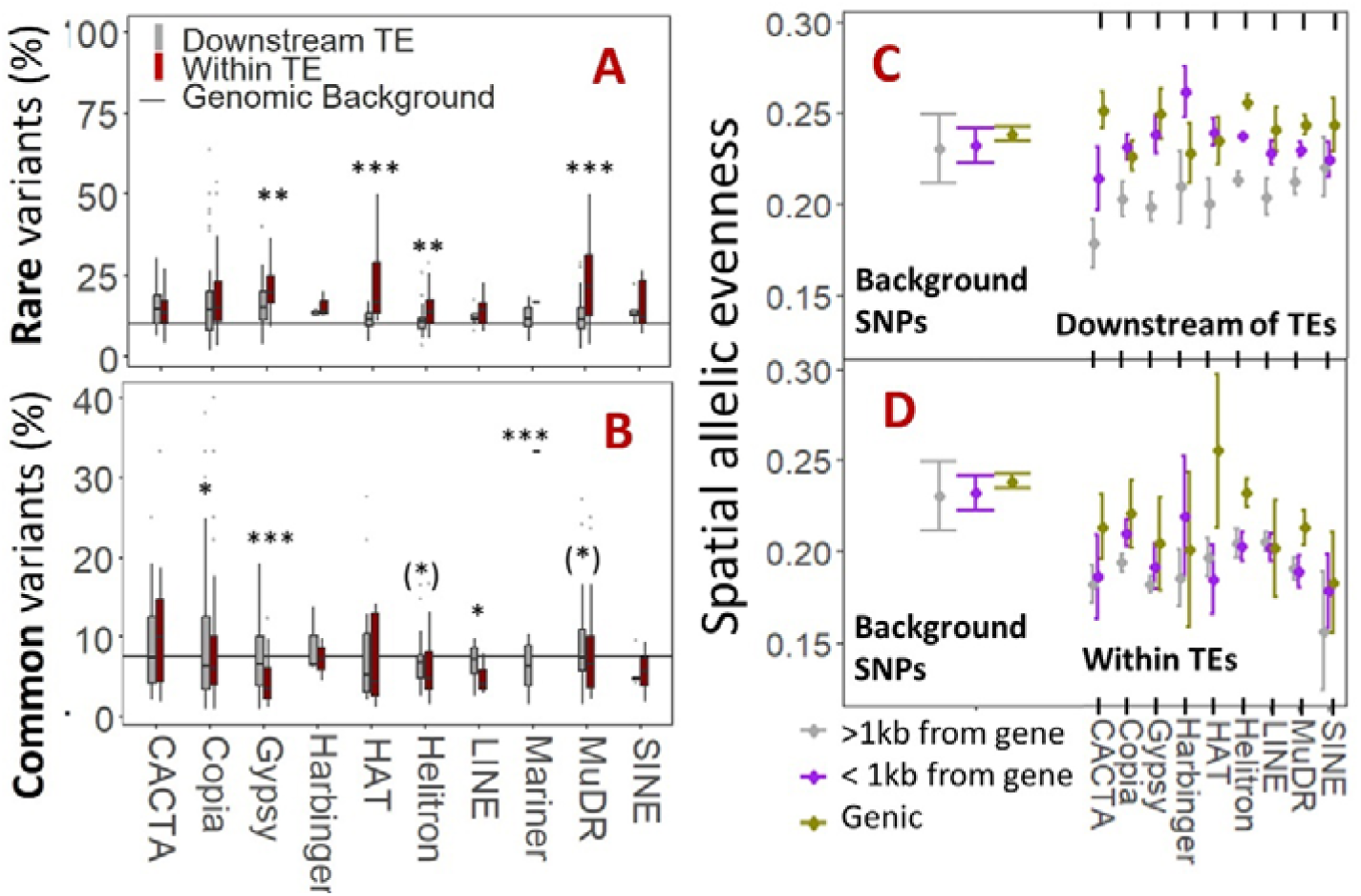
Allelic properties of SNPs within and next to TEs. (**A**) Rare variants (maf between 0.05 and 0.06) were enriched within TEs, (**B**) while common variants (maf > 0.4) occurred more frequent in the genomic regions flanking TEs. (**C**) SNPs downstream of TEs have similar spatial allelic distribution as background SNPs when these TEs are within or near genes, suggesting that SNPs arising near genic TEs are subject to similar evolutionary forces as SNPs far away from TEs. (**D**) On the other hand, high spatial structuring (low evenness) was observed for variants downstream of non-genic TEs and within TEs, which are thought to represent non-deleterious variants that de-activate TEs (Quesneville 2020), or have little impact on their transcription (Lanciano & Cristofari 2020). See Table S5, S6 and S7 for statistical model output.

### Evolutionary signatures downstream of TEs as compared to genomic background

To test whether SNPs downstream of TEs are frequent targets of selection, according to our third hypothesis, we first estimated the tendency of SNPs to be involved in balancing selection based on a spatial nearest-neighbor approach. A lack of spatial genetic structure can be interpreted as a signature of balancing selection (Fijarczyk & Babik 2015, Llaurens et al. 2017). Thus, a high probability of a SNP being different between two nearest neighbor samples across the entire range was taken as a potential signature of balancing selection. For 68,778 SNPs with less than 5% missing data (18419, 44889 and 5358 SNPs within, downstream and away from TEs, resp.), we found that the tendency of SNPs to have balanced spatial distributions is, in general, higher for background SNPs than for SNPs within or downstream of TEs (Fig. 4C and 4D, Table S4). However, some TE superfamilies (CACTA, Harbinger, Helitron) were characterized by increased downstream signatures of balancing selection when residing within or next to genes (high spatial allelic evenness) (4C, Table 1). In addition to these *superfamily*-wide patterns, genetic variants with significantly increased spatial allelic evenness as compared to background genetic variants were particularly abundant in Helitron (downstream of TEs), Harbinger (downstream of TEs flanking genes), and hAT (within genic TEs) *families* (Fig. 6, Table S8 and S9). The Harbinger superfamily is the only superfamily lacking families with significantly reduced spatial allelic evenness (Fig. 6).

The tendency of genetic variants near TEs to be involved in adaptive evolution was assessed through their association with five climatic variables (isothermality, temperature annual range, precipitation seasonality, precipitation of warmest quarter and precipitation of coldest quarter) with pairwise correlation coefficients ranging between 0.03 and 0.42 (Pearson, Table S3). The proportion of SNPs manifesting signatures of diversifying selection as obtained from a climate-informed redundancy analysis (RDA) was highest for background genetic variants (5.3%), followed by SNPs within TEs (3.2%) and SNPs directly downstream of TEs (2.9%) (Fig. 5, Table S4). In accordance, only a few TE superfamilies contained families with significantly increased involvement in climate adaptation (Fig. 6). Interestingly, the finding that background genomic variants away from TEs are more likely to be involved in adaptive processes corresponds to the enrichment of adaptive processes away from TEs (e.g. response to stimuli, and flower, seed and root development) (Table S2). While Mariner, CACTA, and Copia elements frequently manifest significantly enhanced signatures of climate adaptation as compared to background genetic variants (Fig. 6), SINE, LINE, and Helitron elements are nearly completely absent from genomic regions involved in climate adaptation (Fig. 6, Table 1).

**Fig. 5.**
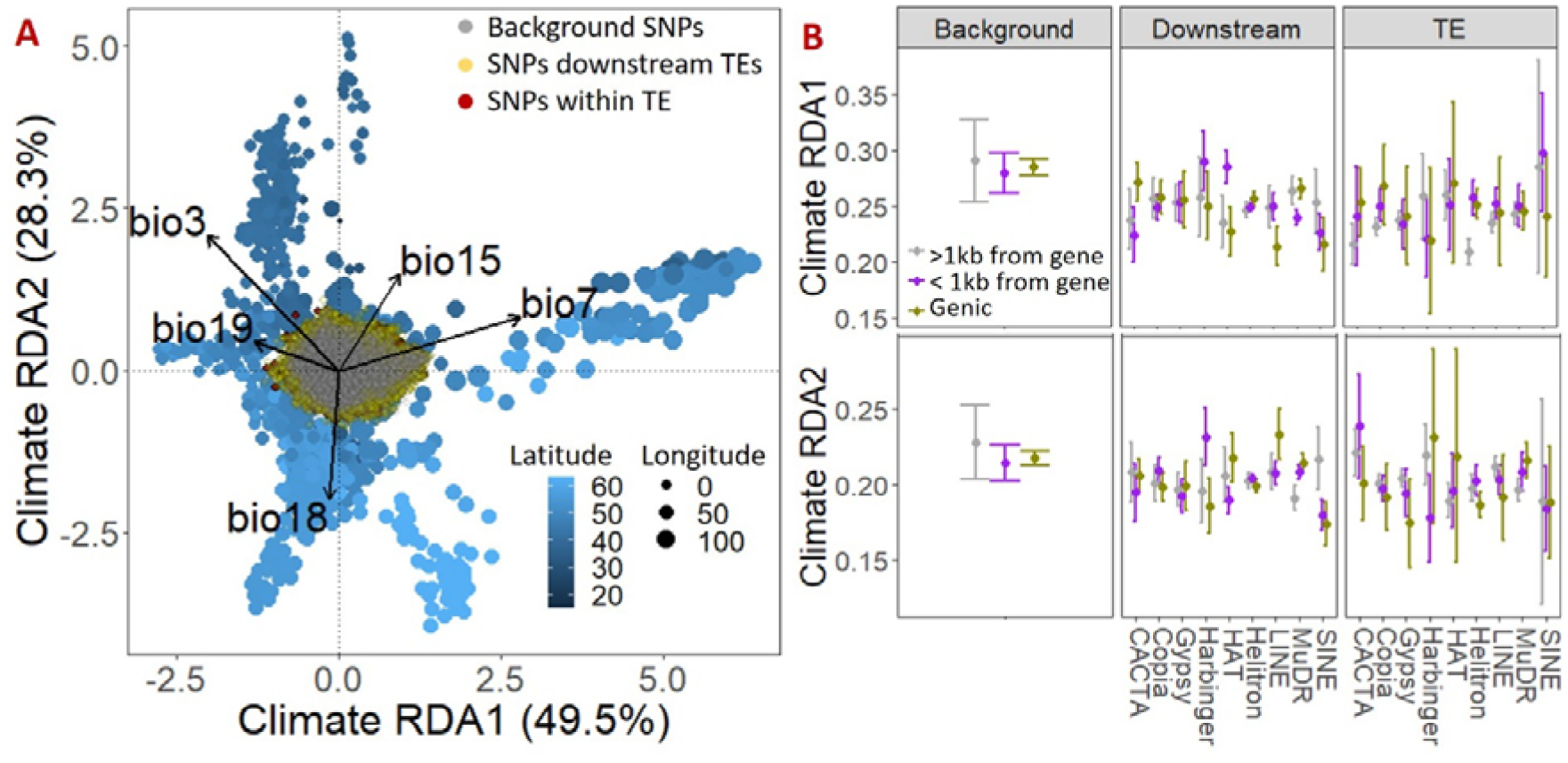
Signatures of climate adaptation within TEs, directly downstream of TE (<1kb) and away from TEs (background) in *Arabidopsis thaliana*. (**A**) A redundancy analysis (RDA) was used to extract outlier SNPs associated with bio 3 (Isothermality), bio 7 (Temperature annual range), bio 15 (Precipitation seasonality), bio 18 Precipitation of warmest quarter and bio 19 (Precipitation of coldest quarter). Alleles mainly correlated with bio7, bio19 (RDA1) and bio18 (RDA2), with higher values of RDA1 representing more continental climates and higher values of RDA2 representing dryer climates. The absolute values of individual SNPs along the RDA axes (RDA loadings; Table S4) represent the strength of their association with climate, and thus the tendency to be involved in selection. (**B**) Background SNPs were more likely than SNPs within or flanking TEs to be involved in climate adaptation.

**Fig. 6.**
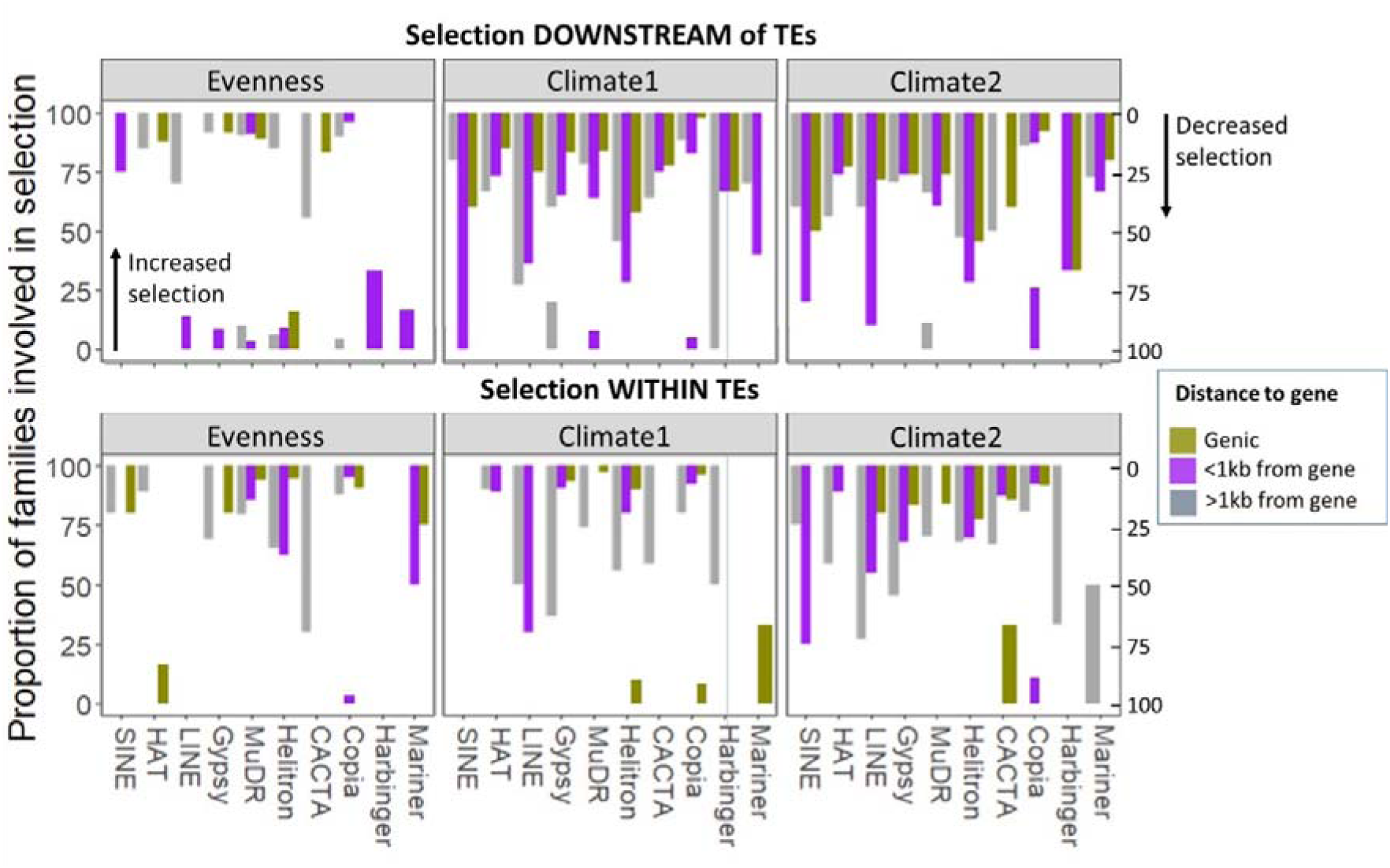
Tendency of TE families to be significantly more or less involved in selection as compared to background genomic variation. , depending on their proximity to genes (Table S8 and S9). Most TE superfamilies are characterized by significantly decreased signatures of selection, climate-related adaptive evolution in particular. However, over 15% of Harbinger and Mariner families flanking genes, and of hAT families within genes, is characterized by significantly increased signatures of balancing selection.

## Discussion

Self-fertilizing species are thought to manifest limited opportunities for evolution: standing genetic variation is expected to be low due to reduced recombination rates, intense inbreeding, and efficient purifying selection. In such species, new mutations may thus play disproportional roles in evolution, particularly when arising non-randomly in functional genomic regions. Here, we demonstrate extreme accumulation of genetic variants within, but also in the direct vicinity of transposable elements (TEs), many of which are enriched near genes with particular molecular or functional processes. In addition, each TE superfamily is characterized by a particular flanking allele frequency spectrum indicative of distinct evolutionary trajectories across superfamilies. The Gypsy superfamily is markedly different from all other superfamilies by its extreme tendency to be underrepresented near genes, particularly those involved in abiotic responses. Together, our results suggest that non-random integration of TEs across the genome combined with high flanking mutation rates, followed by rapid evolution shapes the genome of *Arabidopsis thaliana* and possibly other self-fertilizing species.

Adaptive evolution and stress have frequently been associated with non-random TE mobilization (Cappucci et al. 2019; Baduel et al. 2021), but it remains unclear to what extent this drives non-random mutation as a source for evolution. In *A. thaliana*, where recombination is limited, and nearly exclusive self-fertilization rapidly homogenizes the genome, increased mutation rates in functional genomic regions could generate important opportunities for evolution. We observed marked functional compartmentalization of TE integration across the genome, with several TE superfamilies being significantly over- or underrepresented for important molecular processes. Harbinger elements were exceptionally overrepresented in genes regulating mRNA cis-splicing (at least 5 of all known 31 genes), thus likely impacting transcriptional activity across a considerable part of the genome. Maintaining high genetic diversity in key transcriptional processes across the genome thus appears to be a major evolutionary contribution of Harbinger TEs. Strikingly, the affinity of Harbinger elements to cis-mRNA splicing genes has also been observed in the closely related *A. lyrata* species (De Kort et al. 2022), where genetic variants downstream of Harbinger elements were characterized by balancing selection. The preferential integration of TEs in specific genes of independent study systems indicates that TE distributions are not merely directionless but at least in part predictable.

In addition, regulation of double fertilization, which is key to successful reproduction in all flowering plants, was markedly enriched for TEs of three superfamilies, including Helitron, hAT and MuDR repeats. At least 36 out of 124 genes of the unexplored ECA1 (Early Culture Abundant 1) gametogenesis-related cysteine-rich proteins are flanked by, or contain, TEs. Deregulation of genes involved in fertilization, for example through polyploidization, can induce apomixis (Grimanelli et al. 2001). TEs offer an alternative route towards fertilization gene re-configuration, through local duplication, reshuffling, and transcriptional changes of fertilization genes. Because *A. thaliana* is a ruderal weed frequently characterized by fast reproduction and limited mate availability, genomic mechanisms allowing unsuccessful fertilization to result in uniparental inheritance could offer a powerful evolutionary strategy. Genes specifically expressed in gametes and synergids (such as ECA1) have correspondingly been shown to manifest higher rates of protein evolution compared with genome-wide averages (Gossmann et al. 2014). Although ECA1 has been directly implicated in apomixis in rice (Vernet et al. 2022), it remains unclear to what extent the involvement of TEs and linked genetic variation can cause apomixis in Arabidopsis and other self-fertilizing species.

While TEs thus appear to influence fundamental molecular functions, they are also underrepresented in genomic regions with signatures of climate adaptation and in genes involved in stress responses. The Gypsy superfamily in particular avoids genes (Fig. 1) and manifests downstream signatures of purifying selection: low downstream densities of SNPs (Fig. S3), many of which are low-frequency variants (Fig. S1). Interestingly, while our study suggests that climate adaptation is most efficient in genomic regions free of TEs, TE activity has been frequently associated with stress responses and adaptive evolution (Rey et al. 2016; Schrader and Schmitz 2019; Baduel et al. 2021). Assuming that the 1000 bp region down- and upstream of each TE (with a TE block considered as one collective TE) is directly influenced by TEs, at least 37% of the genome is affected by TEs, rendering it likely that many TEs are associated with selective sweeps. We accordingly identified a few TE superfamilies with a slight (Copia) to moderate (Mariner) increased involvement in adaptive evolution as compared to background genomic diversity (Fig. 6, Table 1). Nevertheless, our direct comparison between TE-flanking regions and background genetic variants far away from TEs, demonstrates that climate adaptation is predominantly associated with genes free of TEs (Fig. 6). Cell division processes are also under-enriched for TEs (Fig. 2), suggesting enhanced molecular protection and/or purifying selection against TE insertion events in vital genomic regions.

The non-random functional distribution of TEs across the genome implies non-random distribution of genetic variants associated with TEs. As a consequence, non-random integration of TEs is expected to offer important opportunities for host evolution. Our findings corroborate this statement, with Harbinger, hAT and Helitron elements predominantly integrating in gene-rich regions (Fig. 1), many of which being involved in important molecular processes (Fig. 2). Genetic variants near these three superfamilies can also reach high frequencies and spatial allelic evenness, in support of their non-deleterious nature and a role in providing a level of evolutionary potential that is beyond what is possible through recombination in a highly self-fertilizing species. In addition, most TE superfamilies are flanked by an allele frequency spectrum that deviates considerably from intra-TE genetic patterns (Fig. S1), suggesting that genomic regions adjacent to TEs are involved in evolutionary trajectories that differ from TE evolution. Most likely, TEs rapidly accumulate neutral genetic variants in line with their high diversification rates (Wicker et al. 2016; Quesneville 2020), while flanking genetic variants caused by mutation-by-insertion, mutation-by-excision, and mutation-by-methylation (Wicker et al. 2016; Habig et al. 2021, Herpin et al. 2021) can provide targets for selection.

The accumulation of SNPs and the distinct allelic patterns flanking TEs point to extensive yet poorly explored opportunities for rapid adaptive evolution in populations lacking a sustainable reservoir of standing genetic variation that can be exchanged between genetically distinct individuals. We argue that, similar to intense outcrossing between populations inhabiting distinct environments, aggregation of genetic variation near TEs can swamp adaptive alleles. This could explain our finding that genomic signatures of climate adaptation are concentrated in TE-free regions of the genome, where adaptive signatures are less subject to mutation. More broadly, where TEs provide genomic redundancy through gene duplication and mutation (cfr. ECA1 gene complex), major opportunities arise for polygenic evolution that cannot be captured by gene-environment association analyses. Thus, where abrupt or alternating environmental changes challenge evolution in the absence of recombination, TE activation and subsequent targeted mutation rate elevation, represent a potent mechanism toward evolutionary rescue.

While our main aim was to explore and identify genetic patterns associated with non-random TE distributions, another major step forward is to unravel to what extent mutations arise through TE insertion, excision or methylation. Because our work is based on TE distributions as determined in the *A. thaliana* reference genome, it is most likely that non-reference TEs contribute to our genome-wide patterns of allelic diversity. As a consequence, our findings are conservative and lack the resolution to disentangle mutations arising from erroneous DNA repair following TE mobilization from mutation-by-methylation. Recent studies point to both mobilization (Baduel et al. 2021) and methylation (Monroe et al. 2022) as major drivers of evolutionary novelty, but how these processes ultimately and interactively shape allelic patterns and evolution across the tree of life remains vague.

## Conclusions

Our results reveal that transposable elements drive the evolution of important biological processes such as double fertilization and transcription in a superfamily-specific fashion, through their non-random genomic distribution. The non-random distribution of TEs is inherently associated with non-random mutation and evolution, as confirmed by allelic signatures deviating significantly from those of background genomic variants. In particular where TEs increase genomic redundancy through gene duplication and subsequent mutation, they offer essential opportunities for rapid polygenic evolution in the absence of recombination. In a broader perspective, our findings incite to consider Lamarckian inheritance of non-random mutations as an important evolutionary strategy in species with reduced recombination, the latter facilitating natural selection based on Darwinian or Mendelian inheritance. Because non-random mutations in functional genomic regions and away from vital functions are less likely to negatively impact plant fitness, we finally argue that non-random mutations arising near co-opted TEs not only allow sustaining high evolutionary rates in self-fertilizing species, but also counteract inbreeding depression through minimizing the deleterious genetic load.

## Methods

### Distribution of TEs and genetic variants

A list of reference TEs (Col0 genome) and their genomic locations was obtained from The Arabidopsis Information Resource (TAIR) (Table S1). Because TEs tend to cluster together, but it remains unknown to what extent TE superfamilies differ in their tendency to be part of TE clusters, we identified TE blocks where at least two TEs were <1kb from each other, and distinguished between solitary TEs (> 5kb), isolated TEs (> 1kb) and block TEs (Fig. 1).

Genetic variants were obtained from the 1001 genomes project (Alonso-Blanco et al. 2016; 1001genomes.org). After filtering on maf > 0.05, a total of 500581, 328665 and 8654 SNPs were found within TEs, within 1 kb downstream of TEs, and away from TEs (> 5kb; background SNPs), respectively. Because mutation rates have been shown to be elevated in the 1000 bp flanking regions of TEs (Wicker et al. 2016), SNPs downstream of TEs were considered in the 1000 bp directly adjacent to any TE. For computational reasons, and because downstream TE regions have been shown to be more likely to be involved in evolution (De Kort et al. 2022), we excluded the 1000 bp upstream regions. We further excluded genomic regions between TEs within blocks, where TEs are < 1000 bp apart because these regions likely or influenced by multiple TE superfamilies both up- and downstream, causing noise in our ability to distinguish TE superfamily-specific effects.

A gene ontology enrichment test was performed to determine whether genes containing TEs are over- or under-enriched for particular molecular processes. We employed the TAIR gene ontology enrichment tool developed for efficient analysis of gene functions in Arabidopsis thaliana, using the Fisher’s Exact test with Bonferroni correction for multiple testing.

### Allele frequency distribution

SNPs with missing data in more than 48 individuals (i.e. 5%) were removed for the analysis of allele frequencies and selection, and missing data imputed with the most common genotype (Table S4). Rare alleles can point to recent mutations, or to deleterious mutations under purifying selection and with limited contribution to evolution. Within transposons, mutations can accumulate rapidly with little impact on nearby genome function (Lanciano and Mirouze 2018; Quesneville 2020), and rare alleles may correspondingly reflect relatively recent, (near-)neutral mutations. Adjacent to transposons, alleles may be rare due to purifying selection or because they are recent and provide opportunities for evolution (evolutionary potential). In self-fertilizing species, new mutations can rapidly shift to fixation locally due to drift, inbreeding and selection, but due to lack of outcrossing, they are not expected to rapidly become common across the range. Common alleles thus represent ancient mutations unless the respective mutations arise repeatedly across the range as a consequence of increased nucleotide sensitivity to mutation (cfr. non-random mutations, Ossowski et al. 2010; Monroe et al. 2022). Common alleles are unlikely to be strongly deleterious and thus likely contribute to the evolutionary potential within the species.

The frequency of rare and common alleles (maf <0.06 and > 0.4, resp.) within and downstream of TEs was statistically compared between TE superfamilies using generalized linear models with Poisson distribution. We expected a high proportion of rare alleles within TEs, particularly in MuDR transposons which are known for their extreme mutation rates (Dupeyron et al. 2019). The proportion of common variants was expected to be higher downstream of TEs than within TEs for many superfamilies, where evolutionary processes contribute to the spatial distribution of alleles.

### Signatures of selection: spatial allelic evenness and climate adaptation

Because existing methods to reveal balancing selection are applicable to the population level (e.g. F_ST_ and SFS approaches), and the samples used here are not organized into populations, we focused on the spatial distribution of alleles as a proxy for the tendency to be involved in balancing selection. Since balancing selection refers to the evolutionary processes of maintaining genetic variation (Fijarczyk and Babik 2015; Han 2019), high genetic dissimilarity between nearby individuals, corresponding to high spatial allelic evenness, may correspondingly point to balancing selection. While gene flow can also enhance spatial reshuffling of alleles, we focused on genetic variants exceeding the spatial allelic evenness of background genomic variation. Specifically, we used a spatial nearest neighbor (NN) approach to study the spatial organization of alleles. The proportion of neighboring individuals with diverging alleles ranged from 0% (alleles are highly spatially clustered) to 56% (alleles are more evenly distributed across space). Spatial allelic evenness was statistically compared between TE superfamilies and background genomic variation, and depending on proximity to nearest gene using generalized mixed models (negative binomial distributions) accounting for random variation due to genomic linkage between TEs within TE blocks (R package “glmmTMB”). We expected variants within TEs to be little involved in balancing selection, and thus to manifest low spatial allelic evenness that do not exceed background allelic evenness. Variants downstream of TEs, particularly those in the vicinity of genes, on the other hand, may frequently engage in balancing selection, resulting in levels of spatial allelic evenness exceeding those of background genomic variation.

The tendency of SNPs near TEs to be more involved in climate adaptation than background genomic variants was assessed through gene-environment association analysis. Specifically, redundancy analysis (RDA) was employed to identify SNPs that align with climate variables (Capblancq and Forester 2021). A total of five bioclim variables downloaded from WorldClim.org (isothermality, temperature annual range, precipitation seasonality, precipitation of warmest quarter and precipitation of coldest quarter) and with low Pearson correlation coefficients (Table S3) were used as explanatory variables, and could be summarized into two orthogonal RDA climate axes significantly associated with genetic variation (Table S3). The absolute values of each SNP onto these climate axes (SNP loadings of “RDA1” and “RDA2”) reflect the tendency of these SNPs to be involved in climate adaptation. Thus, instead of using arbitrary significance thresholds for distinguish outliers from neutral SNPs, we allowed genomic redundancy and polygenic adaptation to contribute to evolution. The tendency of variants to be involved in climate adaptation was statistically compared between TE superfamilies and background genomic variation, and depending on proximity to nearest gene using general mixed models accounting for random variation due to genomic linkage between TEs within TE blocks (R package “glmmTMB”). We expected climate adaptation to be particularly pronounced for genetic variants downstream of TEs near and within genes, given the non-random distribution of TEs in functional genomic regions (Baduel et al. 2021; De Kort et al. 2022).

Because selection may be disconnected from TEs at the superfamily level, we finally used simple linear models to compare signatures of selection of individual TE families to background genomic variants, depending on proximity to nearest gene (within gene, <1kb from gene, >1kb from gene). We then counted the proportion of TE families within each superfamily with signatures of selection exceeding those of background genomic variants.

## Declarations

### Ethics approval and consent to participate

Not applicable

### Consent for publication

Not applicable

## Availability of data and materials

Our data were downloaded from Arabidopsis.org (transposable elements) and from 1001Genomes.org (SNPs). Subsets of these data used for this study are provided as supporting tables.

## Competing interests

The authors declare that they have no competing interests.

## Funding

This work was funded by Research Foundation – Flanders (FWO 12P6521N).

## Authors’ contributions

HDK developed the study, analyzed the data, and wrote the first version of the manuscript. YG performed the high-throughput calculations to obtain genomic distances between transposable elements and nearest genomic variants. FVN supervised the study and commented on the text.

## Acknowledgements

Not applicable.

## Notes

### Competing Interest Statement

The authors have declared no competing interest.

