## Supplementary material for "Non-random polymorphisms near transposable elements facilitate evolution across the genome of *Arabidopsis thaliana*": Fig. S

Supporting Figures

**Fig. S1**. Rare vs. common genetic variants within and downstream of TE superfamilies.

**Fig. S2.** SNP density downstream vs. within TEs.

**Fig. S3.** The probability of a nucleotide to be polymorphic along the 1000 bp downstream a TE, per TE superfamily.

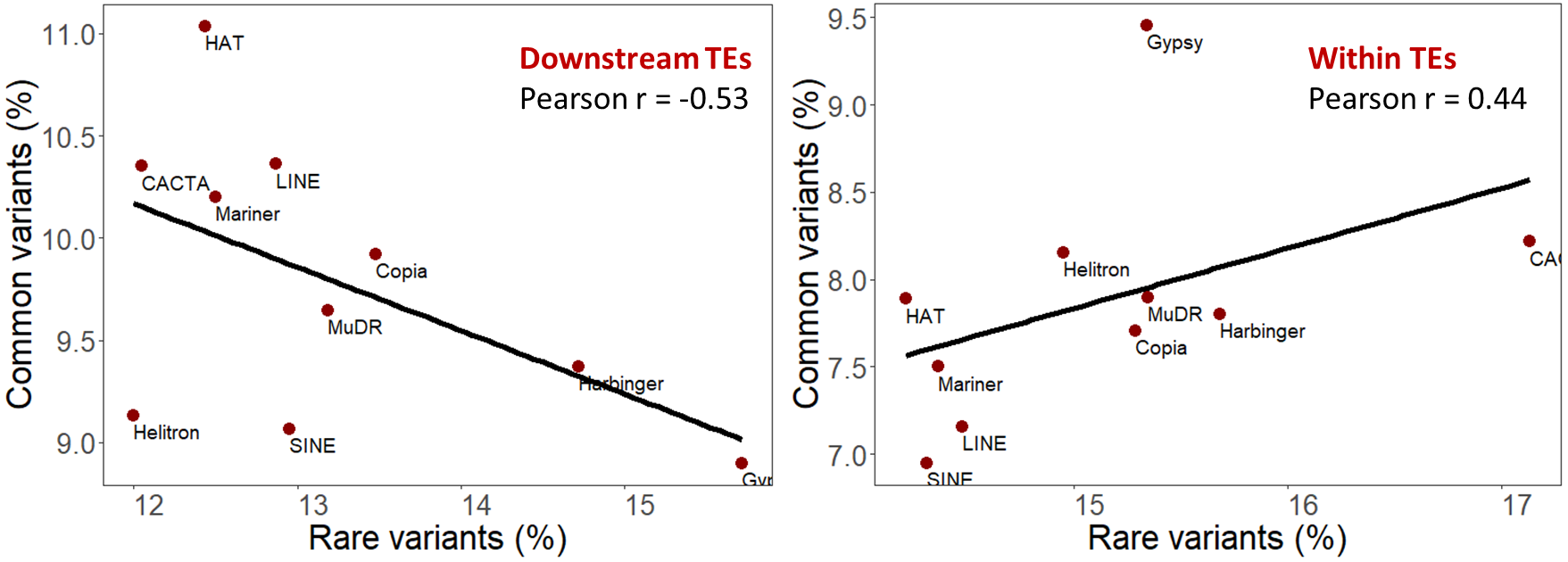

**Fig. S1**. Average allele frequency behavior downstream of, and within, each TE superfamily. Common variants are enriched *downstream of* hAT, CACTA, LINE and Mariner families, while rare variants are enriched *downstream of* Harbinger and Gypsy families, and *within* CACTA elements. Note that the amount of common variants correlates positively with the proportion of rare variants within TEs (right panel), suggesting that difference evolutionary forces shape allele frequency spectra flanking vs. within TEs.

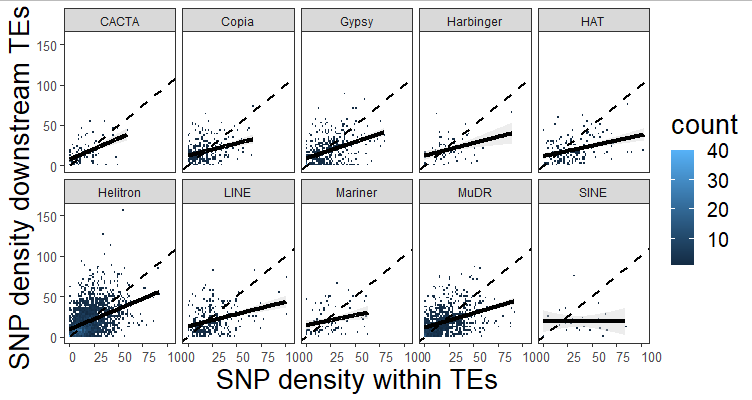

**Fig. S2**. SNP density downstream vs. within TEs. A steeper slope points to a higher accumulation of SNPs downstream of TEs relative to within TEs.

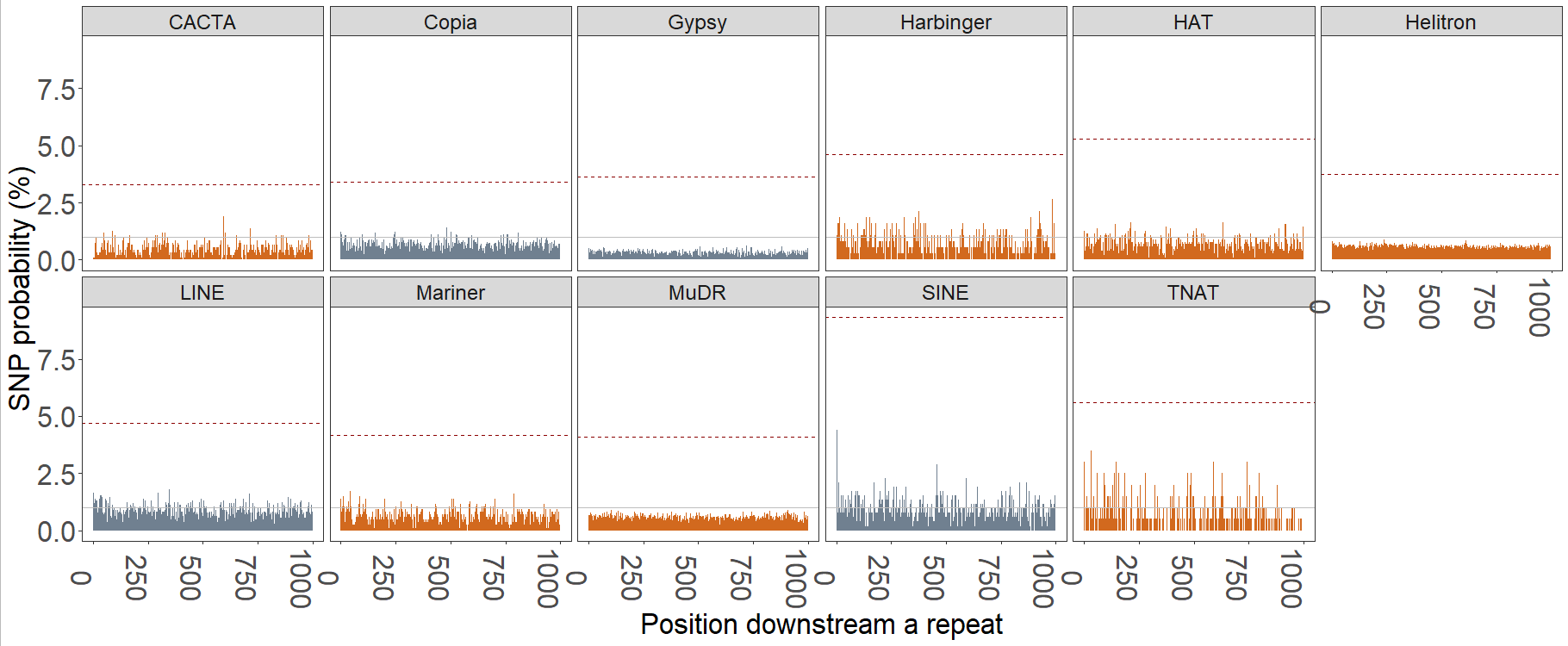

**Fig. S3**. The probability of a nucleotide to be polymorphic along the 1000 bp downstream a TE, per TE superfamily. Where the SNP probability crosses the 1% horizontal line, there is a chance of 1 in 100 to encounter a SNP. The dashed red line represents the SNP probability in the TE center (20bp central region), and is generally much higher than the downstream SNP probability, suggesting that genetic variants within TEs can accumulate without strong deleterious effects on the organism. In general, the highest probability for genetic variants downstream of TEs is found near TNAT elements.

**On average, a SNP with maf > 0.05 occurred every 47 bp within and every 51 bp downstream of TEs (*i.e.* 21.3 SNPs and 19.5 SNPs per 1kb, resp., Supp. Fig. x, Table Sx). Moreover, the probability of encountering a SNP in the 1kb region directly flanking TEs decreased only slowly with distance from TE, and is similar among superfamilies (Supp. Fig. x). Although extreme SNP peaks can be found in the TE centers (Supp. Fig. x), the average SNP density within TEs is not higher than in the 1kb downstream region (Supp. Fig. x). On the contrary, particularly where high SNP densities are reached within and adjacent to TEs (*e.g.* for SINE elements, Fig. appendix), the gravity center of SNPs lies downstream of TEs (Fig. 3A). Genetic variants thus tend to accumulate downstream of TEs (Fig. 3A, Supp. Fig. x) particularly in the case of TE superfamilies enriched for gene-flanking repeats (Fig. 3B, Fig. 3C). The SINE superfamily, for example, can accumulate extreme downstream SNP densities and is characterized by a relatively high proportion of repeats integrating next to genes (Fig. 3B, Fig. 3C).**
